# Validity of the Lumen^®^ hand-held metabolic device to measure fuel utilization in healthy young adults

**DOI:** 10.1101/2020.05.05.078980

**Authors:** Kent A. Lorenz, Shlomo Yeshurun, Richard Aziz, Julissa Ortiz-Delatorre, James R. Bagley, Merav Mor, Marialice Kern

## Abstract

**Background:** Metabolic carts measure the carbon dioxide produced and oxygen consumed from the breath in order to assess metabolic fuel usage (carbohydrates vs. fats). However, these systems are expensive, time-consuming, and only available in the clinic. A small hand-held device capable of measuring metabolic fuel via CO_2_ from exhaled air has been developed

**Objective:** To evaluate the validity of a novel hand-held device (Lumen^®^) for measuring metabolic fuel utilization in healthy young adults

**Methods:** Metabolic fuel usage was assessed in healthy participants (n = 33; age: 23.1 ± 3.9 y) via respiratory exchange ratio (RER) values from the “gold-standard” metabolic cart as well as %CO_2_ from the Lumen device. Measurements were performed at rest in two conditions, fasting, and after consuming 150 grams of glucose in order to determine changes in metabolic fuel. Reduced major axis regression was performed as well as Bland-Altman plots and linear regressions to test for agreement between RER and Lumen %CO_2_.

**Results:** Both RER and Lumen %CO_2_ significantly increased after glucose intake compared with fasting conditions (*P* < .0001). Regression analyses and Bland-Altman plots revealed an agreement between the two measurements with a systematic bias resulting from the nature of the different units.

**Conclusions:** This study shows the validity of Lumen^®^ to estimate metabolic fuel utilization in a comparable manner with the “gold-standard” metabolic cart, providing the ability for real-time metabolic information for users under any circumstances.

## Introduction

Respiratory quotient (RQ) is the ratio of carbon dioxide produced to oxygen consumed (VCO_2_/VO_2_) measured directly at the cellular level to estimate metabolic fuel utilization. However, this method requires the insertion of a catheter into the vein and artery for a blood sample or taking a tissue sample, which makes the RQ measurement invasive and infeasible outside of laboratory setting [1]. With the use of a metabolic cart, an indirect measure of RQ can be made by indirect calorimetry through respiratory gas exchange of O_2_ and CO_2_. This is the respiratory exchange ratio (RER), which is currently the preferred method for determining metabolic fuel utilization. Unlike RQ, RER measures the carbon dioxide produced (VCO_2_) and oxygen consumed (VO_2_) from exhaled air [1,2]. Both RQ and RER indicate the relative contribution of carbohydrate and lipid to energy expenditure [3]. Though RER measurement is not invasive, this method is time-consuming (up to 40 minutes) and is only available in test laboratory setting and it requires technical and physiological expertise for handling the metabolic cart and interpretation of the metabolic data obtained.

Metaflow Ltd. developed Lumen^®^, a novel metabolic fuel utilization breathalyzer which is a personalized hand-held device that provides an individuals’ metabolic state in real-time, by measuring CO_2_ from exhaled breath (Figure 1). The device indirectly measures metabolic fuel usage via a CO_2_ sensor and a flow sensor, to determine the rate of CO_2_ production from a single breath maneuver. The CO_2_ concentration in the exhaled volume of air is determined from a specific breathing maneuver with a breath hold of 10 seconds. This concept is based on the fact that oxygen consumption is stable under resting conditions [4]; thus, a change in the metabolic fuel use will generally be represented by changes in CO_2_ production. For carbohydrate oxidation more carbon dioxide is produced, relative to the consumption of oxygen. For fat oxidation, less carbon dioxide is produced. Thus, from the resulting RER, for a resting state ranging between 0.7 and 1, the contribution of lipid or carbohydrate usage can be determined. The breath maneuver enables quantifying the changes in CO_2_ production, allowing the user to estimate their metabolic state [5]. The use of a smartphone application enables the user to track metabolic status outside of physiologic test laboratories.

**Figure 1.**
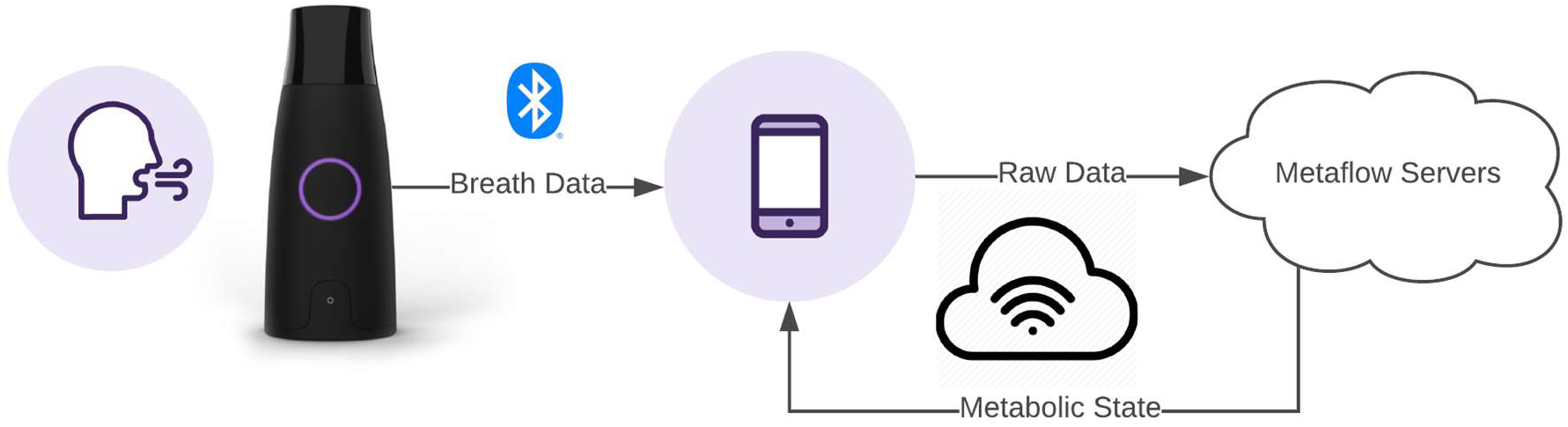
A schematic representation of the Lumen^®^ device and application.

Internal studies (unpublished data) showed the potential of Lumen to accurately measure the metabolic fuel usage in response to diet and exercise, comparable to the “gold standard” metabolic cart. In this study, we aim to evaluate agreement between Lumen measurement and metabolic cart in healthy participants before and after glucose ingestion under stable resting conditions.

## Methods

### Participants

Fifty-four healthy volunteers reported to the Exercise Physiology Laboratory from the Department of Kinesiology at San Francisco State University to participate in this study. To be included for the study, participants must have been between the ages of 18-45 years old; with a BMI less than 30 kg/m^2^;and not doing any high intensity aerobic exercise training over 3 times per week and without any known cardiovascular-, pulmonary-, and/or metabolic diseases. The study was approved by the University’s Institutional Review Board for Human Subjects, and written informed consent was obtained from each participant before testing.

### Study Design

Participants were recruited and their height and weight measured using a stadiometer and Seca scale (Seca, Hamburg, Germany). If they met the BMI criteria, they were provided their own Lumen device which was labeled with their unique identification number. The Lumen device was paired and synchronized to the participant smartphone together with the Lumen application. Participants practiced the Lumen breathing technique while supervised and took the device home for further familiarization period in order to show proficiency with the device and application. They were instructed to perform Lumen metabolic measurements for at least 30 sessions, with each session consisting of 3 breath maneuvers, and to take 3 sessions at different time points each day. After the minimum amount of home breath sessions were collected, participants were scheduled for the study laboratory measurement day. All participants came to the test laboratory between 07:00 a.m. and 11:00 a.m. after a 12-h fast and had abstained from any form of physical activity (other than walking).

On the laboratory testing day, blood glucose samples were taken by sterile finger prick blood sample and measured by a glucometer (OneTouch, LifeScan Inc. Milpitas, CA). For the indirect calorimetry measurement, the participant had to lay down in supine position on a padded examination table, where a rigid clear plastic canopy with a comfortable, flexible seal was placed over the head and upper part of the torso. Once the metabolic cart measurement was ended, the participant was seated in a comfortable chair. After five minutes of rest, they were asked to perform two Lumen breath sessions (5-minute break between each session). A valid Lumen session measurement for the evaluation with a difference of less than 0.2 %CO_2_ between the breaths in a session. The first Lumen session immediately after the metabolic cart measurement was used for data analysis. In case with an invalid first session, the second session was analyzed.

Once finished, participants were asked to drink 150 grams of a glucose solution (3 servings of 50 grams with 20 minutes intervals between each serving). Forty-five minutes after the intake of the first drink (corresponding to 5 minutes after finishing the last serving), their glucose levels were reassessed, and the same assessment procedures as during the fasted state before the glucose intake were repeated. Participants were removed from the analysis if they were unable to finish all glucose drinks.

### Metabolic cart

RER was analyzed using a calibrated TrueOne^®^ 2400 metabolic cart (ParvoMedics, Murray, UT, USA). This system uses a paramagnetic oxygen analyzer and infrared carbon dioxide analyzer with a Hans Rudolph heated pneumotach. The ParvoMedics system was warmed up for at least 60 minutes each day before testing, in order to ensure accurate and stable readings. The gas analyzers and flow sensor were calibrated as per manufacturer’s recommendations. Calibration of the analyzers performed using a high precision gas mixture (O_2_, CO_2_, remainder N_2_) and calibrated and accepted with a less than 0.1% error with the calibration gas. Flow and volume were calibrated using a calibrated 3 L syringe (Hans Rudolph, model 5530) to ≤ 1% error. The ambient temperature was kept between 22 and 26°C in the test laboratory. Relative humidity was maintained stable at roughly 60%. Once calibration was acceptable and complete, a ventilated hood with subject cover was placed over the participant’s head and positioned around the upper torso area to ensure no escaping air from the hood. The participants were required to stay awake during the measurement procedure. The hood ventilation (VE) was measured during the recording, and CO_2_ and O_2_ concentrations were measured from it. VCO_2_, and VO_2_ parameters were calculated and taken as 30-s averages. For this study we defined the subject steady-state metabolic measurement based on observed variations in the VO_2_ and VCO_2_ of less than ≤ 5% CV for period of at least five consecutive minutes. Inability to meet these criteria resulted in removal of the data from the analysis.

### Lumen

During the measurement day, participants took 2 sessions of 3 Lumen breaths after the metabolic cart. The Lumen breathing maneuver consists of three phases, starting from the end of a normal expiration (functional residual capacity). The participant takes a deep breath in through the Lumen device, followed by a 10 second breath hold. Afterwards, the subject exhales through the Lumen device, with a steady exhalation flow to at least the starting level of the maneuver. The Lumen smart phone application guides the participant through each phase of the lumen maneuver. Each Lumen session was repeated after a 5-minute pause interval. Validity of breath maneuvers was systematically evaluated by the Lumen application. Inability to perform valid Lumen breath measures resulted in removal of the data from the analysis.

### Statistical Analyses

All variables were tested and visualized for normal distribution before the tests.

To evaluate the changes after glucose intake, two-tailed paired parametric t-tests were performed for blood glucose levels, RER levels, and Lumen %CO_2_ before and after glucose intake.

For agreement validations, major axis regression (‘Deming’s method’) was performed in order to compare RER of the metabolic cart and %CO_2_ from the Lumen device [6]. As RER and %CO_2_ are in different units, the analysis is identical to ordinary least product regression (also known as reduced major axis regression), which is the most suitable analysis for comparison between two methods of measurement [7]. Moreover, Bland-Altman plot was created to demonstrate limits of agreement, together with a linear regression to test the distribution of error to identify if any systematic bias existed between Lumen and RER measures. In addition, a simple linear regression (ordinary least squares) was performed to determine the ability to predict Lumen values from the gold-standard value of RER.

Statistical analyses were performed using GraphPad Prism 8 (GraphPad Software Inc., LA Jolla, CA). The threshold for significance was set at *P* < .05.

## Results

From the original fifty-four participants recruited, twelve were excluded prior to laboratory testing and nine had to be excluded during the testing day for failing to meet the inclusion criteria as detailed in the methods section: one participant was unable to consume all glucose drinks due to nausea, three participants did not achieve 5 mins of stable metabolic cart measurement (CV < 5% in VO_2_ and VCO_2_), and five participants were unable to perform a valid Lumen measurement (Figure 2). Characteristics of the final thirty-three participants presented in Table 1.

**Table 1.**
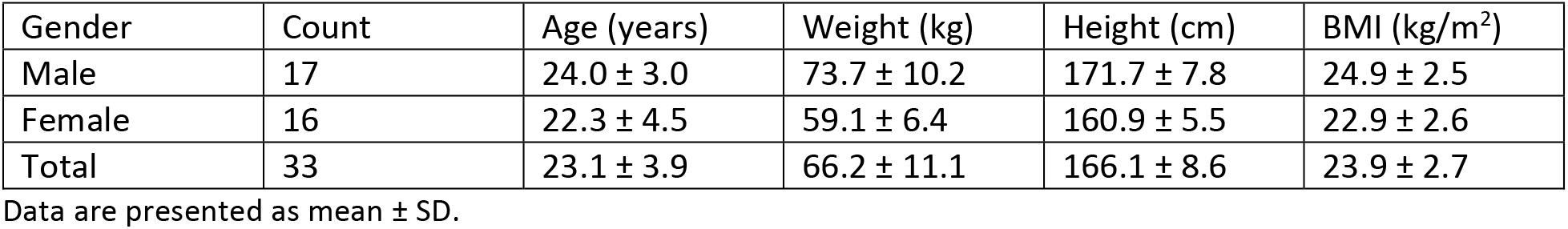
Descriptive statistics of study’s participants.

| Gender | Count | Age (years) | Weight (kg) | Height (cm) | BMI (kg/m<sup>2</sup>) |
| --- | --- | --- | --- | --- | --- |
| Male | 17 | 24.0 ± 3.0 | 73.7 ± 10.2 | 171.7 ± 7.8 | 24.9 ± 2.5 |
| Female | 16 | 22.3 ± 4.5 | 59.1 ± 6.4 | 160.9 ± 5.5 | 22.9 ± 2.6 |
| Total | 33 | 23.1 ± 3.9 | 66.2 ± 11.1 | 166.1 ± 8.6 | 23.9 ± 2.7 |

Data are presented as mean ± SD.

**Figure 2.**
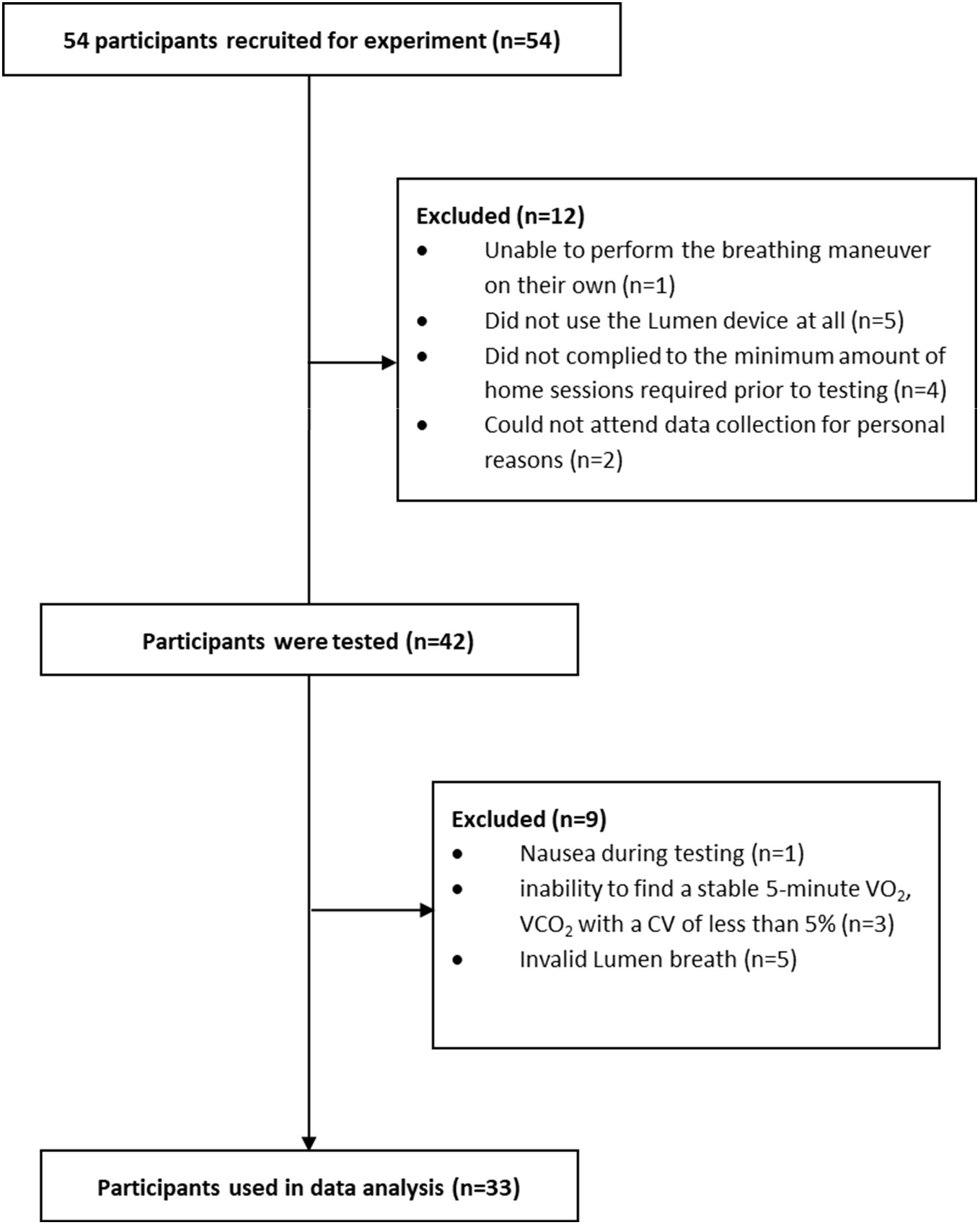
Consolidated Standards of Reporting Trials (CONSORT) flow diagram.

Blood glucose levels increased from 90.6 ± 9.2 mg/dL to 145.2 ± 25.3 mg/dL as a result of glucose intake (*t*(32) = 11.04, *P* < .0001; Figure 3A). RER levels increased from 0.787 ± 0.043 to 0.876 ± 0.053 in response to glucose intake (*t*(32) = 10.84, *P* < .0001; Figure 3B). Moreover, Lumen %CO_2_ concentrations significantly rose from 4.20 ± 0.4 to 4.48 ± 0.34 (*t*(32) = 5.978, *P* < .0001; Figure 3C). These analyses have confirmed the ability of both the metabolic cart and Lumen to detect changes in metabolic fuel utilization.

**Figure 3.**
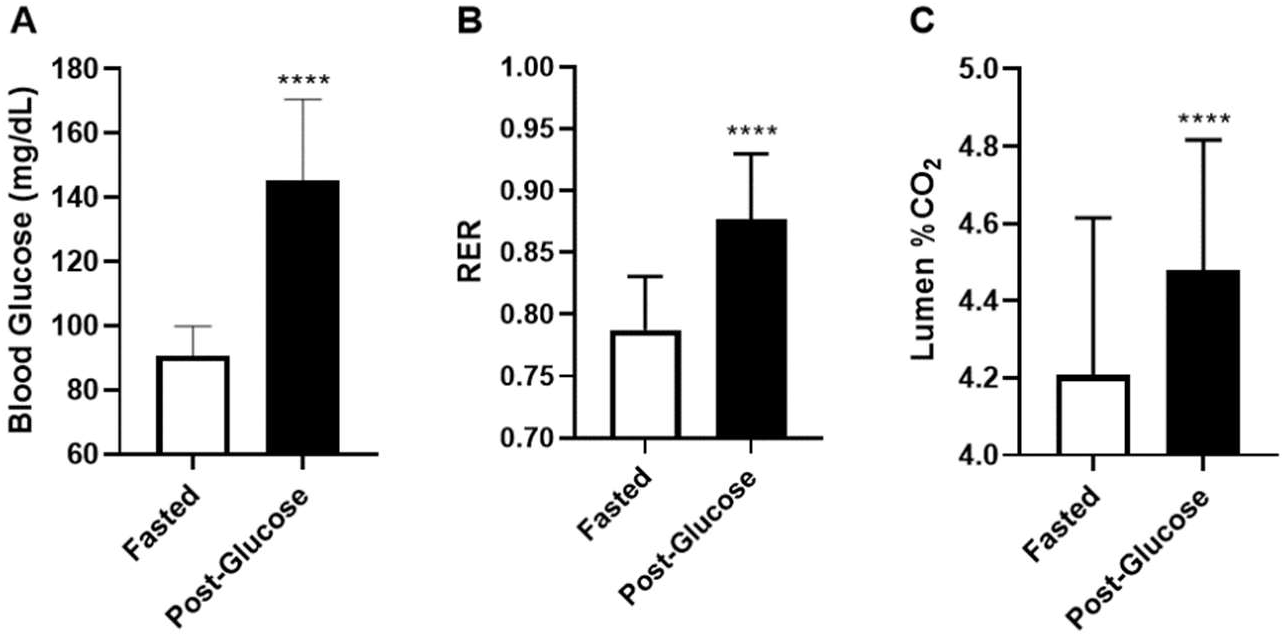
Changes of blood glucose (A), RER (B), Lumen %CO_2_ (C), after glucose intake. Data are presented as mean ± SD. \*\*\*\**p* < 0.0001. *n* = 33 for each state.

In order to look for agreement between RER units from the metabolic cart and %CO_2_ from Lumen, reduced major axis regression was performed [8]. It revealed a significant relationship between RER and Lumen %CO_2_ (*F*(1,63) = 18.54, *P* < .0001, y = 6.111x - 0.7445, x-intercept = 0.1218; Figure 4). This analysis validated the agreement between Lumen %CO_2_ and metabolic cart RER, with a systemic bias resulting from the nature of the different units.

**Figure 4.**
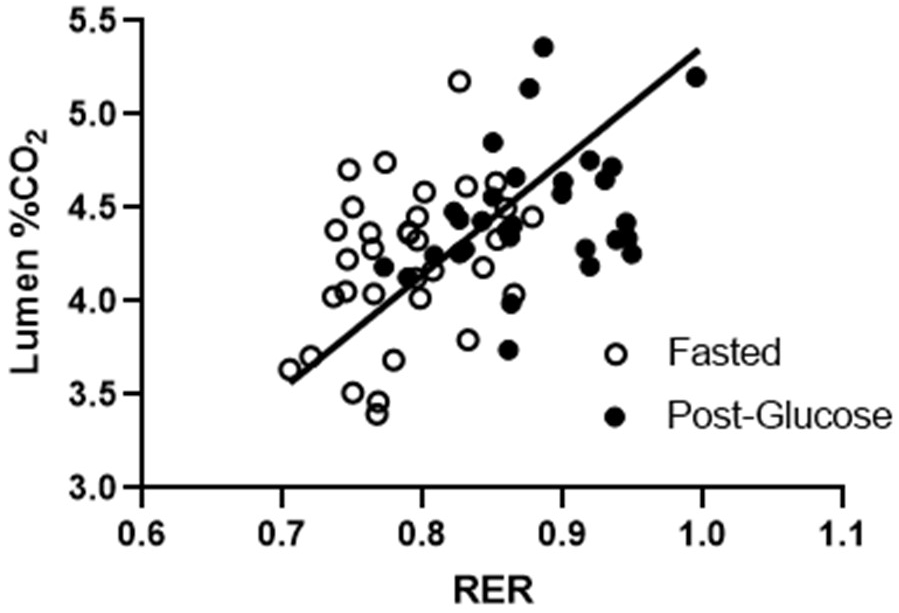
Reduced major axis regression of RER from the metabolic cart and Lumen’s %CO_2_ measurements for metabolic activity. *n* = 33 for each state.

A Bland-Altman plot was made to discuss limit of association between RER and Lumen %CO_2_ [9]. Bland-Altman plots are used to calculate the level of agreements between two measures by studying the mean difference between measurements and constructing limits of agreement [10]. Bland-Altman analysis revealed a mean difference between RER units and Lumen %CO_2_ of 3.505 with 95% limits of agreement between 2.784 to 4.226 (Figure 5). Since the Lumen device always provides a measurement that is numerically much higher than RER, Lumen measurements with smaller values should produce smaller differences and larger Lumen values will produce larger differences. To test the bias in measurements, a regression of difference (Lumen %CO_2_ – RER) scores on average ((Lumen %CO_2_ + RER)/2) values was performed. Simple linear regression showed that a significant model effect of the bias (*F*(1,63) = 721, *P* < .0001, *R*^2^ = 0.9196). As the vertical distribution of scores are tight to the line, it indicates that the errors are consistent across a range of values.

**Figure 5.**
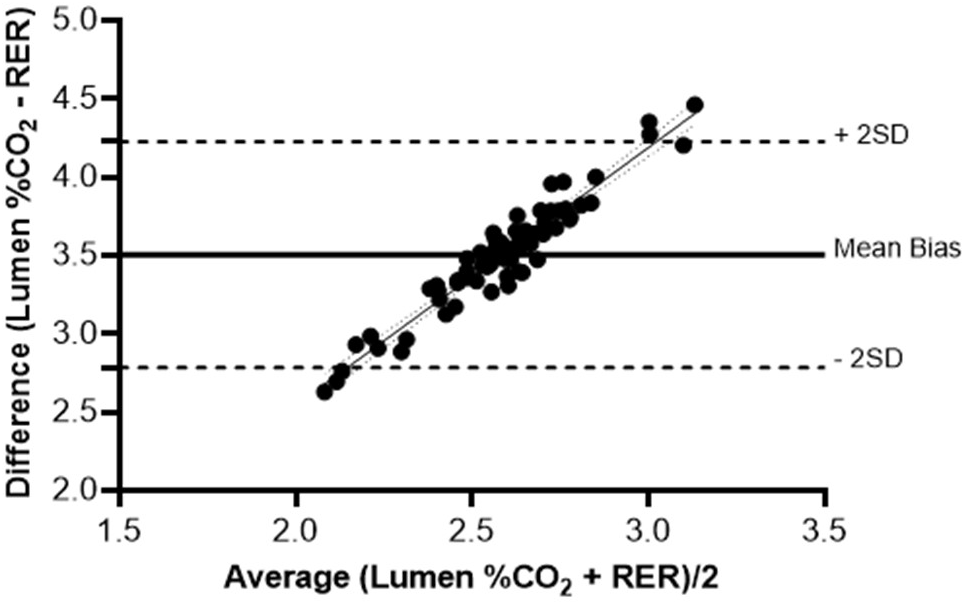
Bland-Altman analysis with simple linear regression. The solid line represents the mean bias between the Lumen %CO_2_ and RER. The upper and lower dashed lines represent the 95% confidence intervals (±2 SD) from the mean bias. Linear regression line: y = 1.646*x - 0.7472. *n* = 33 for each state.

To determine the ability of RER to predict Lumen %CO_2_, ordinary least squared regression was performed to estimate Lumen values from RER measures, with the assumption that RER is an accurate measure. With RER as the independent variable, we used linear regression to predict Lumen %CO_2_, and a significant model effect was present (*F*(1,63) = 18.54, *P* < .0001, *R*^2^ = 0.2274; Figure 6). The RER parameter estimate indicated that for every one-unit increase in RER, a 2.914-unit increase (*SE* = 0.6767) in Lumen %CO_2_ is expected. However, since a full unit increase in RER is not a plausible outcome, this parameter estimate can be interpreted similarly by saying a 0.1-unit increase in RER (e.g., 0.7 to 0.8) will produce a 0.2914-unit increase in Lumen %CO_2_.

**Figure 6.**
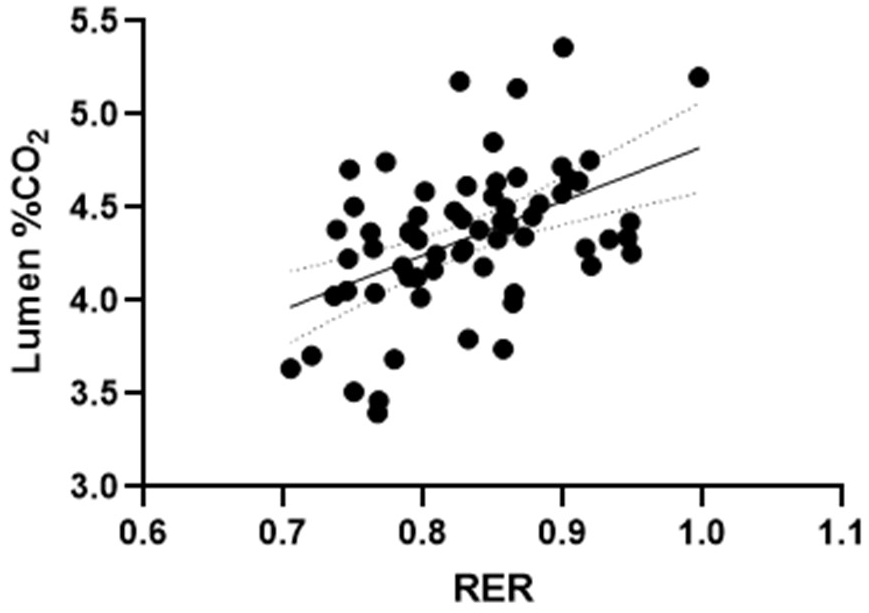
Ordinary least squared regression of RER and Lumen %CO_2_. *n* = 33 for each state.

## Discussion

This study evaluated the capability of the Lumen device to assess changes in the body’s metabolic fuel utilization, compared to gold standard indirect calorimetry metabolic cart measurement in healthy young adults. Our results show that Lumen %CO_2_ levels are in agreement with RER values from the metabolic cart, which correspond to relative changes in metabolic fuel utilization.

Both Lumen %CO_2_ and metabolic cart RER showed significant increases in metabolic levels, as a result of glucose intake in healthy individuals in resting conditions (Figure 3). These results can be expected as cells using more carbohydrates as fuel, produce more CO_2_ relative to the O_2_ consumption compared to cells metabolizing fat. The ratio between the CO_2_ production and the O_2_ consumption in this process is known as respiratory quotient (RQ) or RER. RQ and RER vary depending on the energy source of the cell (carbohydrate vs. fat), and the acronyms are commonly used interchangeably [1–3]. In resting conditions, oxygen consumption is fairly stable [11,12], meaning that participants’ changes in the RQ are due to changes in CO_2_ production. This is the concept underlying the Lumen device, enabling it to track changes in metabolic fuel utilization. For that reason, it was important to ensure that participants in this study were at rest before and during their measurements.

Reduced major axis regression revealed an agreement between RER and Lumen %CO_2_ (Figure 4). This analysis enables to test for agreement between methods with different units and verified the validity of the Lumen device with the gold standard metabolic cart. The Bland-Altman plot showed a mean bias of 3.505, and when used in conjunction with a significant linear model of difference scores compared to mean values [13], support the agreement between the two methods despite measuring in different units (Figure 5). These results demonstrate the ability of Lumen to provide comparable results to the metabolic cart in assessing the metabolic fuel utilization.

Furthermore, the results from the simple linear regression predicting Lumen %CO_2_ using RER values, suggest that while there is measurement agreement between the Lumen %CO_2_ and RER, the proportion of explained variance remains low (Figure 6). Thus, Lumen can be seen to be an effective instrument for monitoring relative, individual changes in metabolic responses (within-subject consistency), rather than a substitute for the metabolic cart (between-subject precision).

Evidence suggests that the assessment of RER can be a benefit for multiple conditions, such as nourishing, diabetes prevention, weight management, physical activity, and healthy lifestyle [14,15]. It has previously been shown that RER could be a prognostic marker of weight loss and a predictor of weight gain [16,17]. Moreover, minute-by-minute RER corresponded directly to intensity of exercise, and slopes of RER were different in response to different dietary treatments [18]. However, although RER is currently the preferred method for determining metabolic fuel, it is a costly, time consuming, uncomfortable and an impractical tool for assessing metabolic activity over time and for real time day to day usage. In contrast, the Lumen device is small, relatively cheap, mobile, user-specific, delivers the outcome immediately to the user and enables taking real time decisions.

### Limitations

This study is the first to show agreement between Lumen %CO_2_ and RER. However, it is important to note that participants in this study were young (mean age: 22.4 y) and healthy individuals. With increasing age, metabolism changes as can be seen in metabolic cart studies [19–21]. Future studies will need to examine whether RER metabolic cart levels correspond to Lumen CO_2_% levels in older subjects and those with metabolic conditions.

Another limitation is that unlike the metabolic cart, the Lumen device does not measure oxygen consumption. Accordingly, the Lumen measurement should be performed under resting conditions with subsequent stable VO_2_, allowing the correct interpretation of changes of %CO_2_ as changes in metabolic state.

In addition, results from this study show high peak of blood glucose levels 45 minutes after glucose intake (5 minutes after the 3^rd^ drink), whereas both RER and Lumen %CO_2_ showed a more moderate increase in levels. Therefore, it is possible that the metabolic cart and Lumen measurements were performed too early, as it may be that in some of our participants the peak glucose occurred later than at 45 minutes, thus not yet fully metabolized [22].

## Conclusions

In summary, Lumen^®^ can provide valid information regarding an individual's resting metabolic state, and in agreement with results from metabolic cart. Unlike the metabolic cart, Lumen measurement can be performed anywhere, without the need for a specialized lab, equipment, and technical support. The capability of taking metabolic measurements continuously can provide numerous insights about the metabolic state of an individual, for further scientific knowledge and understanding about metabolism and nutrition.

## Conflict of Interest

SY and MM are employees of Metaflow Ltd., and contributed to the design and analysis of the study as well as the preparation of the manuscript. The other authors declare no conflicts of interest.

## Ethics Statement

This study was approved by the University’s Institutional Review Board for Human Subjects, and written informed consent was obtained from each participant before testing.

## Author Contributions

KAL analyzed the data and prepared the manuscript SY analyzed the data and prepared the manuscript RA coordinated the project and collected the data JO coordinated the project and collected the data JRB reviewed and edited the manuscript MM conceived, designed, and supervised the study as well as reviewed and edited the manuscript MK conceived, designed, and supervised the study as well as reviewed and edited the manuscript All authors approved the manuscript before submission

## Funding

This work was supported by Metaflow Ltd.

## Acknowledgements

We would like to thank the participants for their time in taking part in this study, and the Lumen team for their support. We would like to acknowledge Casey Curl for his work in setting up the protocols and procedures during preliminary testing.

